# Impact of the nucleosome histone core on the structure and dynamics of DNA containing pyrimidine-pyrimidone (6-4) photoproduct

**DOI:** 10.1101/2020.04.24.060012

**Authors:** Eva Matoušková, Emmanuelle Bignon, Victor Claerbout, Tomáš Dršata, Natacha Gillet, Antonio Monari, Elise Dumont, Filip Lankaš

**Affiliations:** Department of Informatics and Chemistry, University of Chemistry and Technology Prague, Technická 5, 166 28 Prague, Czech Republic; Université de Lyon, ENS de Lyon, CNRS UMR 5182, Université Claude Bernard Lyon 1, Laboratoire de Chimie, F69342, Lyon, France; Université de Lorraine and CNRS, LPCT UMR 7019, F-54000 Nancy, France; Institut Universitaire de France, 5 rue Descartes, 75005 Paris, France

## Abstract

The pyrimidine-pyrimidone (6-4) photoproduct (64-PP) is an important photoinduced DNA lesion, which constitutes a mutational signature for melanoma. The structural impact of 64-PP on DNA complexed with compaction proteins, and notably histones, affects the mechanism of its mutagenicity and repair but remains poorly understood. Here we investigate the conformational dynamics of DNA containing 64-PP lesions within the nucleosome core particle by atomic-resolution molecular dynamics simulations at the multi-microsecond time scale. We demonstrate that the histone core exerts important mechanical restraints that largely decrease global DNA structural fluctuations. However, we also show that local DNA flexibility at the damaged site is enhanced, due to imperfect structural adaptation to restraints imposed by the histone core. In particular, if 64-PP faces the histone core and is therefore not directly accessible by the repair protein, the complementary strand facing the solvent exhibits higher flexibility than the corresponding strand in a naked, undamaged DNA. This may serve as an initial recognition signal for repair. Our simulations also pinpoint the structural role of proximal residues from the truncated histone tails.

## INTRODUCTION

The UV-induced (6-4) pyrimidine-pyrimidone photoproduct (64-PP, Figure 1A) is the second most frequent DNA photolesion after cyclobutane pyrimidine dimer (CPD) and is commonly produced as a result of type II photosensitization. Although 64-PP is much more efficiently repaired than CPD, and hence is more readily eliminated from the cells (1–3), it also has a much higher degree of mutagenicity (4–6) and as such it is correlated to the development of malignant skin cancer (7). From the chemical point of view, it involves a single covalent bond linking the pyrimidine (thymine) and pyrimidone moieties (denoted by Thy and Pyo, respectively, Figure 1A), as opposed to the formation of two covalent bonds in CPD. This chemical difference is also correlated to an increase in the global flexibility of the lesion, which leads to complex structural signatures also due to the rearrangement of the DNA double helix to accommodate the highly distorted dimer. The increased flexibility and the structural distortion may also explain the higher repair rate of 64-PP vs. CPD (8,9).

**Figure 1.**
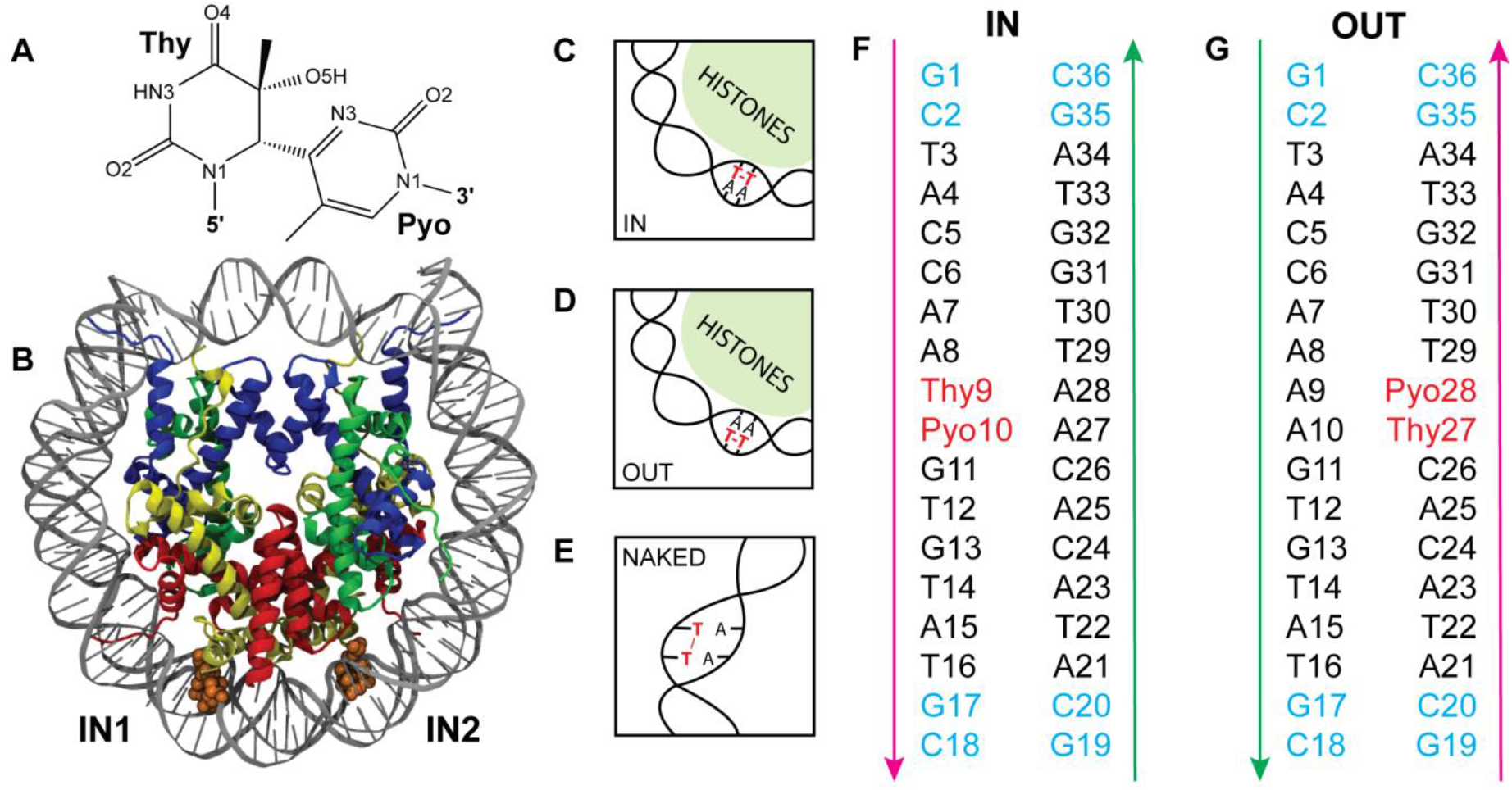
(A) Chemical structure of the 64-PP lesion. The thymine moiety is denoted by Thy, Pyo refers to the pyrimidone. The lesion arises as two adjacent thymines in the same DNA strand are linked by a single covalent bond between their C6 and C4 atoms. (B) Crystal structure (4YM5) used as a starting point for the MD simulations. The two symmetrically positioned lesions (orange) face the histone core (IN). The H2A histone chain is rendered in yellow; H2B in red; H3 in blue; H4 in green. The structure is schematically represented in (C). Besides that, an analogous structure (4YM6) with the two lesions in the complementary strand and therefore facing the solvent (OUT) was also investigated (D), together with the naked, damaged (E) and control undamaged B-DNA oligomers where Thy and Pyo were each replaced by a thymine. In the naked simulations, the nucleosome DNA sequences (black, with the lesion in red) were capped with GC dinucleotide steps (blue) to avoid end-effects (F, G). Arrows in (F,G) indicate the 5’ to 3’ direction.

Several experimental and computational studies have focused on the determination of the structural dynamics of duplex DNA oligomer containing the 64-PP lesion. Early NMR investigation of short DNA duplexes of the CGCA<u>TT</u>ACGC sequence (underscore denotes the 64-PP position) suggested an overall 44° bend of the double helix (10,11), while a pioneering 800 ps unrestrained molecular dynamics (MD) simulation of the same duplex indicated a much smaller bending of 13.6° (12). A recent MD simulation study of this sequence, embedded in a 16 base pair (bp) DNA duplex and extended up to 2 μs, has evidenced a larger flexibility and a polymorphic character, most notably indicating that both straight and highly bent conformations are accessible (9). Besides that, an experimental work using Förster resonance energy transfer (FRET) on a different sequence (GCATCG<u>TT</u>CTCATA) excluded a highly bent structure but reported substantial unwinding and elongation of the damaged duplex compared to the intact B-DNA. Thus, the effect of 64-PP on the DNA duplex structure most probably depends also on the specific nucleobases surrounding the lesion, pointing to strong sequence effects. Obtaining a specific structural signature of 64-PP is highly desirable, also due to the potential harmfulness of this lesion (7). Furthermore, it has been shown that the specific DNA sequence markedly affects both the extent and accuracy of the translesion DNA synthesis (13). Thus, although the structural features of naked, 64-PP-containing, DNA duplexes are still not fully understood, the current knowledge suggests rich conformational dynamics affected by the sequence context.

However, in a biologically relevant context, DNA will be coiled around compaction proteins, leading to complex architectures. Such constrained environment may significantly alter the DNA structural dynamics, also considering the extremely important mechanical energy related to DNA compaction (14). In eukaryotic organisms, the double-stranded DNA wraps around a core of eight histone proteins (two copies each of H2A, H2B, H3 and H4, see Figure 1B) to constitute the so-called nucleosome core particle (NCP). NCP represents the fundamental unit of the compaction system, the ensemble of NCPs connected by linker DNA gives raise to the chromatin and higher level chromosome organization. Most notably, chromatin is also a highly dynamic system and the change in its compaction level is associated to epigenetic modulations.

A high-resolution structure of NCP has been solved around 20 years ago (15,16) and the nucleosomal organization has been extensively studied using a diverse range of experimental and computational methods, including MD simulations (17–23). However, the impact of such a peculiar macromolecular environment on the structure and dynamics of the damaged DNA in the nucleosome (N-DNA) is a much less addressed question that consequently stays almost completely unanswered, despite the potentially important consequences for repair, transduction, or gene expression. For instance, Greenberg and coworkers have experimentally quantified a 100-fold enhancement of DNA-protein cross-link reactivity for abasic sites in NCPs vs. B-DNA (24).

In the present contribution we aim at tackling this issue and answering the following basic question: To what extent does the structural and dynamical effect of a 64-PP photolesion differ in B-DNA vs. N-DNA? To this end, we rely on all-atom, multi-microsecond molecular dynamic simulations to characterize the structural outcome of 64-PP embedded in a DNA duplex coiled in NCP, profiting from two recent X-ray structures obtained by Osakabe *et al.* (25) (PDB ID codes 4YM5 and 4YM6). Each structure features two occurrences of 64-PP, symmetrically positioned with respect to the NCP dyad axis. In the 4YM5 structure, the photolesions are facing the histone core (denoted as inwards, IN, Figure 1-B, C, F). In contrast, the 4YM6 structure contains two symmetric 64-PP lesions facing the solvent (outwards, OUT, Figure 1D, G).

Importantly, Osakabe and co-workers (25) report that the electron densities around the 64-PP, either in IN or OUT positions, are difficult to resolve in both strands, suggesting that the DNA region including or in close proximity to 64-PP retains higher flexibility even in the nucleosome. This observation leads to the hypothesis that the flexibility around 64-PP might serve as a recognition signal for the nucleotide excision repair (NER) system and in particular for the UV-DDB complex, an initiation factor for the nucleosome excision repair pathway. However, the structural disorder at the 64-PP regions hampers any more detailed structural insight. Hence, extensive atomic-scale molecular dynamics, as performed in our study, is a viable alternative to provide the atomic level description of the structural constrains imposed by the NCP and their subtle interplay with the inherent flexibility and polymorphism of the damaged DNA.

## METHODS

### Molecular dynamics simulations

The OL15 force field (26) was consistently used to describe the DNA, while the histone was described using Amber ff99SB (27) force field, the TIP3P model (28) was chosen for water. The 64-PP was parameterized following the Amber Antechamber procedure compatible with the Amber force field. Charges were obtained from the fitting of the electrostatic potential obtained from quantum chemical calculations using the RESP methodology of Amber. The system was solvated in an octahedral periodic box and neutralized by K^+^ ions parameterized according to Joung and Cheatham (29). For the nucleosome simulations, the crystal structures 4YM5 and 4YM6 were used as starting points. Ten to thirty residues of the H2A and H2B histones tails, close to the damages, are missing in the PDB structures and are absent in our simulations. In the following, the DNA in NCP will be referred to as N-DNA. The naked simulations were started from the canonical B-DNA conformation where the two central thymines were mutated to 64-PP. A standard equilibration protocol consisting of a series of constrained energy minimizations and short MD runs was employed, as detailed in the Supplementary Information. The nucleosome simulations (ca. 200,000 atoms) were extended to 3.5 μs, those of the naked damaged DNA to 20 μs and the control undamaged DNA to 4 μs each. Sequences of 14 base pairs (bp) around the damaged sites in the nucleosomes (Figure 1-F, G) were analyzed. They are denoted IN1, IN2, OUT1, OUT2 for the two instances of the lesion in each NCP structure. In the naked DNA oligomers, the same sequences were further capped by GC dinucleotides at each end to limit end effects, yielding 18-bp oligonucleotides (Figure 1-F, G). The capping GCs were excluded from any analysis. As a control undamaged B-DNA, the same 18-bp sequences were simulated, with two central thymines in place of the 64-PP lesion.

### Data analysis

The DNA MD trajectories were analyzed at various levels of detail, including standard local single-strand base-to-base coordinates as defined in the 3DNA algorithm (30). The most coarse-grained level consists in quantifying the relative displacement and rotation of the two helical regions flanking the damaged site (Figure 2). Base-fixed reference points and frames (31) in four Watson-Crick base pairs at each side of the lesion were averaged to yield effective rigid bodies (32). Their relative displacement and rotation were described using rigid body coordinates exactly analogous to the local base pair step coordinates as defined in the 3DNA algorithm (30). In the following, such global parameters are called interhelical (IH) shift, IH slide and IH rise for translations, IH roll, IH tilt and IH twist for rotations. While the standard base pair step coordinates describe local relative displacement and rotation of the reference points and frames fixed to the two neighbouring pairs, our IH coordinates analogously report the global configuration of the two helical domains flanking the damaged site (Figure 2). In this way it is possible to describe the structural effect of the lesion in the full six-dimensional interhelical space. The rigid body construction and coordinate definitions are detailed in ref. (32).

**Figure 2.**
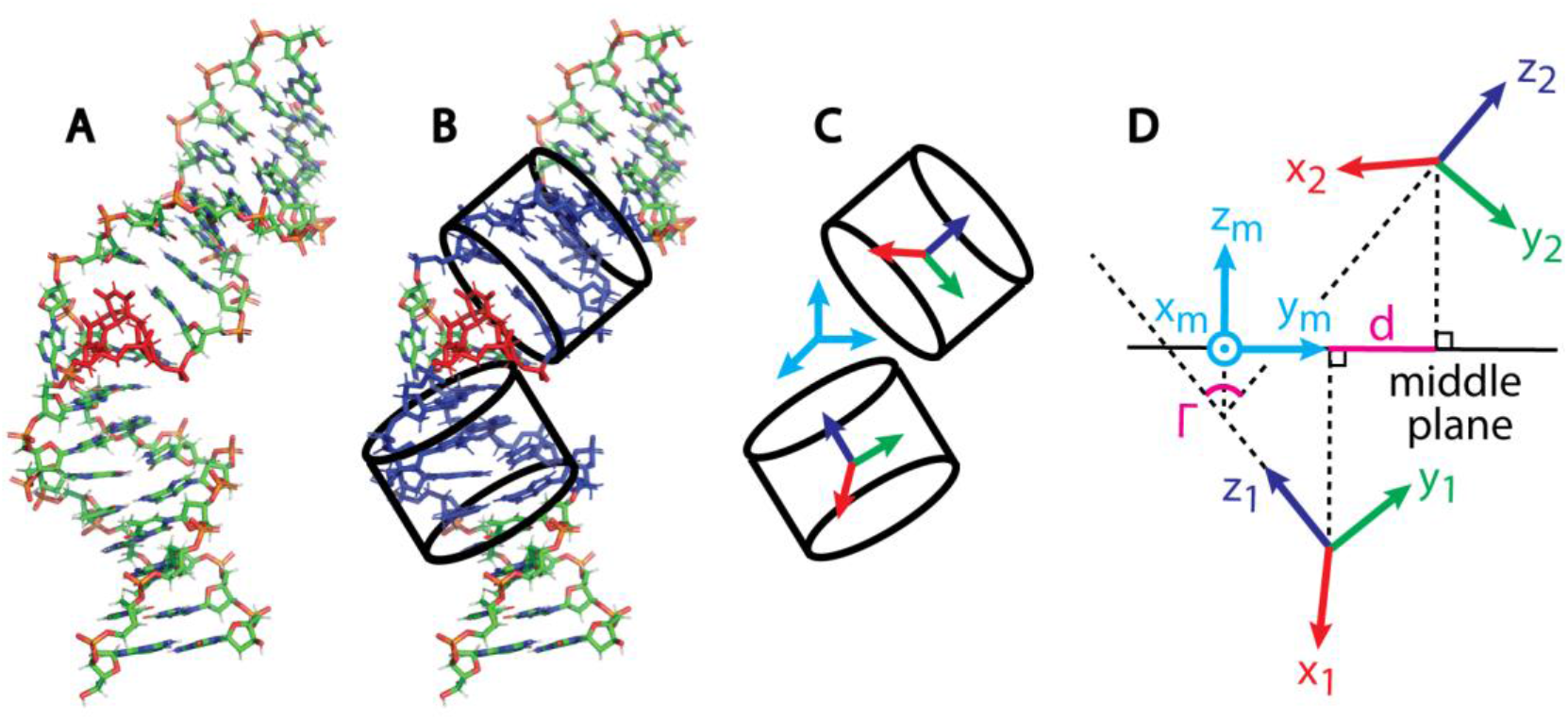
Interhelical coordinates. In each MD snapshot (A), reference points and frames of bases in four pairs (B, blue) on both sides of the 64-PP lesion (B, red) are averaged to obtain mean reference points and frames of effective rigid bodies (C), following the direction of the reference strand. The relative displacement and rotation of the bodies are described by interhelical (IH) coordinates as detailed in the text. To define the IH coordinates, an auxiliary frame (middle frame, cyan in (C) and (D)) is first constructed. Its x-y plane, the middle plane, serves to measure the lateral displacement magnitude d between the rigid body reference points, while the bending magnitude Γ is the angle between the z-vectors of the body frames. The x-axis of the middle frame points towards the major groove at the lesion site, serving to determine the direction of bending and lateral displacement. The damaged strand (magenta arrow in Figure 1F, G) is chosen as the reference strand for IN2 and OUT1 sites, the undamaged one (green arrow) is the reference strand for IN1 and OUT2. This choice ensures consistent definitions of the IH coordinates with respect to the dyadic symmetry of the nucleosome.

The two-component vector (ρ, τ) of IH roll ρ and tilt τ defines the interhelix bending. However, it may be more informative to work in polar coordinates and use an equivalent description in terms of the bending magnitude Γ and bending direction φ, related to roll and tilt as

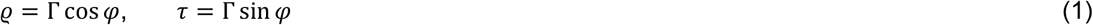

The direction φ = 0 indicates bending towards the major groove, while φ = π (or 180°) means bending towards the minor groove. Similarly, the lateral displacement of the two helices is defined by the two-component vector (ξ, ζ) of IH shift ξ and IH slide ζ. Just as for bending, we introduce the lateral displacement magnitude *d* and direction *Ψ* by the relations

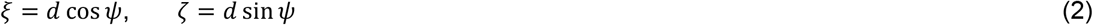

This global description is complemented by local single-strand coordinates capturing displacement and rotation of each individual base relative to the preceding one in the same strand. The base-to-base coordinates obtained for each strand separately may be more informative in our case than the more standard base pair step coordinates, since here the two strands are expected to behave differently depending on whether they bear the lesion or not, or whether they face the histone core or are exposed to the solvent. Assuming that the coordinates of a given strand taken together exhibit a multivariate Gaussian distribution, their entropy is given by (33)

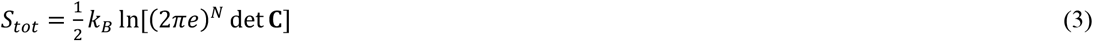

where **C** is the covariance matrix of the coordinates, *N* is the number of coordinates (degrees of freedom), *k_B_* the Boltzmann constant and *e* the base of the natural logarithm. It is convenient to work with entropy per coordinate, defined as

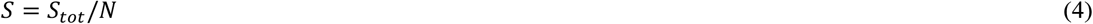

Indeed, while the total entropy is extensive, the entropy per coordinate is an intensive quantity and therefore it allows for direct comparison between systems with different numbers of degrees of freedom.

Finally, hydrogen bond lengths, C1’-C1’ distances and angles were computed and the Cartesian coordinate clustering performed using the *cpptraj* module of Amber. The DPSCAN method (34) was employed for the clustering. The first μs of the naked simulations and the first 0.5 μs of the nucleosomal ones were excluded from the analysis to avoid the effect of the major conformational rearrangements taking place at the beginning of the simulation. Errors are estimated by repeating the calculations for the first half and the second half of the trajectory (or first and second half of its portion belonging to a particular state) and taking the mean difference with respect to the whole trajectory. The DNA conformation in the crystal structure of the DNA-DDB2 complex ((35), pdb code 3EI1) is used for structural comparisons.

## RESULTS

The interhelical (IH) conformational dynamics inferred from the simulations is summarized in Figure 3, Table 1 and Table S1. The first major structural feature differentiating B-DNA and N-DNA is the bending magnitude. We observe in all cases that the N-DNA is bent in the vicinity of 64-PP by 35-57°, whereas the naked sequences exhibit bending angles of at most 20° (see Table 1). Also, the interhelical flexibility is reduced: Figure 3 (left) shows the bending magnitude and direction of the damaged N-DNA in comparison to the naked damaged and control undamaged DNA, analogous data for the lateral displacement are in Figure 3 (right). These data highlight the narrower range of the interhelical fluctuations around the damage in the presence of the histone core: while the 64-PP lesion enhances the flexibility of the B-DNA compared to the undamaged duplex, especially in the IN sequence, the conformational ensemble sampled by the damaged N-DNA is much more restricted. Note that the two damaged sites in the nucleosome are not structurally fully equivalent in the nucleosome crystal because of different crystal packing context (25). This fact may explain the differences seen in the MD data (red and magenta in Figure 3).

**Table 1.** Ensemble averaged interhelical (IH) bending and lateral displacement magnitudes and directions. The quantities are defined and errors are computed as detailed under Methods.

| System | $\Gamma$ (°) | $\varphi$ (°) | $d$ (Å) | $\psi$ (°) |
| --- | --- | --- | --- | --- |
| OUT-naked | $16.95 \pm 0.49$ | $15.39 \pm 1.31$ | $3.21 \pm 0.00$ | $352.1 \pm 0.27$ |
| IN-naked | $15.72 \pm 8.41$ | $335.49 \pm 24.15$ | $2.11 \pm 0.74$ | $274.29 \pm 42.31$ |
| IN-naked, last 6μs | $20.15 \pm 1.96$ | $217.24 \pm 4.4$ | $1.05 \pm 0.19$ | $211.47 \pm 3.94$ |
| IN1 | $42.69 \pm 1.24$ | $62.70 \pm 2.80$ | $0.23 \pm 0.21$ | $9.52 \pm 76.73$ |
| IN2 | $34.82 \pm 1.03$ | $68.77 \pm 1.70$ | $1.40 \pm 0.66$ | $342.35 \pm 7.26$ |
| OUT1 | $39.63 \pm 0.14$ | $129.87 \pm 2.57$ | $4.05 \pm 0.06$ | $313.45 \pm 1.64$ |
| OUT2 – State 1 | $51.41 \pm 0.55$ | $93.36 \pm 2.71$ | $1.53 \pm 0.15$ | $292.11 \pm 14.67$ |
| OUT2 – State 2 | $56.59 \pm 1.04$ | $96.51 \pm 0.78$ | $5.48 \pm 0.00$ | $326.74 \pm 1.05$ |
| Control IN | $7.58 \pm 0.11$ | $235.82 \pm 0.38$ | $0.64 \pm 0.01$ | $8.58 \pm 0.64$ |
| Control OUT | $6.85 \pm 0.01$ | $234.37 \pm 1.06$ | $0.45 \pm 0.00$ | $10.72 \pm 0.79$ |
| DNA-DDB2 crystal | 34.18 | 21.87 | 2.43 | 171.52 |

**Figure 3.**
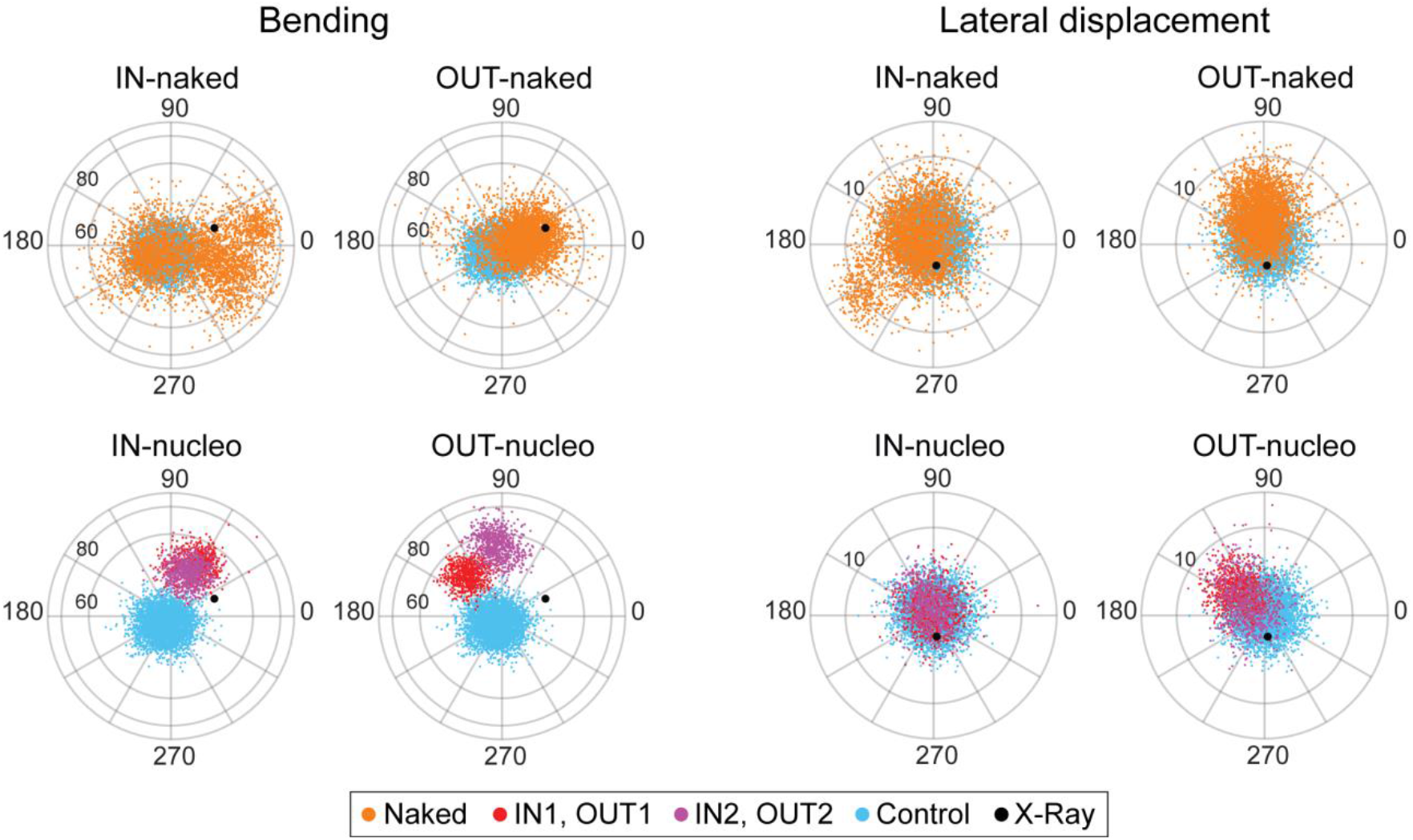
Polar plots indicating the magnitude Γ and the direction φ of bending (left), and the lateral displacement magnitude d and direction Ψ (right) of the two helical fragments flanking the lesion (see Methods and Figure 2 for the definitions). The direction around 0°(or 360°) indicates bending or displacement towards the major groove at the damaged site, values around 180° represent the minor groove direction. The naked, damaged DNA (upper part, orange), and in particular the IN sequence, exhibits a wide range of conformations. These structural fluctuations are substantially reduced in the nucleosome (lower part) where the DNA is also significantly bent. While the two nominally equivalent IN nucleosomal sites (IN1, IN2) behave similarly, important difference is observed between the two OUT sites. The data for the control undamaged B-DNA (cyan) and for the crystal structure of the DNA-DDB2 complex (black dot) are shown for comparison. The bending magnitude is in degrees, the lateral displacement magnitude in angstroms. The ensemble averaged values are in Table 1.

The local dynamics around 64-PP in the naked duplexes is strongly sequence-dependent. Indeed, clustering of the MD data in the Cartesian space (see Methods) shows that the naked OUT duplex is characterized by only one dominant conformation in which 64-PP assumes an extra-helical position (Figure S1). This structure is moderately bent towards the major groove and displaced in a similar direction (Table 1). In contrast, the naked IN duplex explores four markedly different conformational states, each of them lasting for multiple microseconds (Figure S2). Importantly, the conformational states include both nearly straight (clusters 1 and 4 in Figure S2) and bent (clusters 2 and 3) conformations, while 64-PP can assume both intra- and extra-helical positions. In particular, the most populated cluster 4, covering roughly the last 6 μs of the simulation (blue in Figure S2), is moderately bent and displaced but this time towards the minor groove, and the local structure around the lesion is also very different from the OUT case: the C1’-C1’ distances are both around 14 Å and the glycosidic torsions χ of the two adenines facing the lesion are in the *syn* domain.

The inclusion of the NCP profoundly modifies the previous scenario. The two IN sites of the N-DNA now take just one conformation each (Figure S3) and are structurally very similar, exhibiting IH bending of about 30-40° directed roughly half-way between the major groove and the backbone (φ ≈ 60 − 70°) and a slight displacement towards the major groove (Table 1). In contrast, the N-DNA OUT sites, in which 64-PP faces the solvent, display a particular behaviour - our simulation suggests three well-defined conformational states. More specifically, while OUT1 only explores one single conformational state, OUT2 displays two different states, each of them capable to survive for > 1 μs (Figure S4). The OUT1 state is bent by similar amount as the IN states but in a different direction, and is displaced further into the major groove. The two OUT2 states are both severely bent by > 50° in similar directions but differ in the displacement magnitude (Table 1).

The near structural equivalence of the two IN sites and the different geometries of the three conformations adopted by the OUT sites are confirmed by the mean values of the single-strand (base-to-base) local coordinates (Figures S5-S8). The highly distorted geometry of the strand complementary to the lesion and facing the solvent at the IN sites (Figure S6) features an extreme pattern of twist (ca. 60° at the AA inter-base step facing the lesion and 10° in the preceding step) as well as a very high shift (4 Å). Such values are not expected in a nucleosome structure containing intact DNA, where the duplex is only moderately deformed (15,16).

The N-DNA structural states also differ in their flexibility, which we quantify by means of the conformational entropy. The entropy values were computed from the local single-strand, base-to-base translational and rotational coordinates as defined in 3DNA (30) (see Methods for details). Only the 64-PP lesion (or the two bases facing it in the complementary strand) and one more adjacent base at each side were considered. The entropy calculation assumes a multidimensional Gaussian (normal) distribution of the coordinates. We thoroughly verified that, if separated into the conformational states (see above), the individual 1D distributions of the single-strand coordinates involved in the computation are all close to Gaussian, adding confidence to the normality assumption. Notice that the entropy calculation involves the full coordinate covariance matrix, so that possible covariances (couplings) between different coordinates are taken into account.

The entropy values are plotted in Figure 4. At the IN sites, the damaged strand facing the histone core is rigid (has low entropy), as expected. In contrast, the undamaged strand facing the solvent displays higher flexibility around the lesion and is even more flexible than the corresponding strand in the undamaged control oligomer. As for the strands at the OUT sites, the individual states differ in their flexibility, some of them having lower entropy than the control undamaged DNA and others higher. In OUT1 and especially in state 2 of OUT2, the undamaged strand is still flexible (has high entropy), even though it faces the histone core.

**Figure 4.**
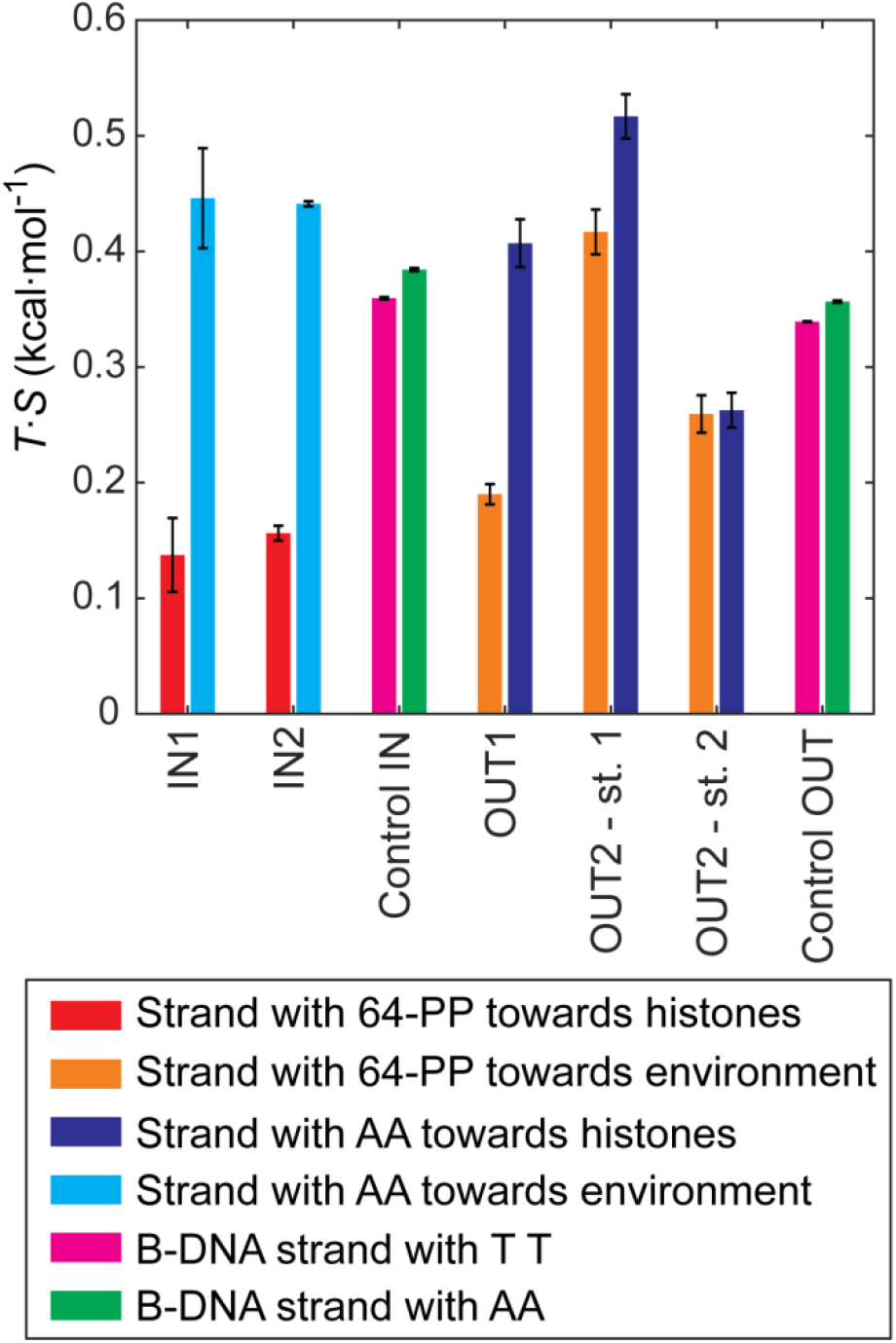
Conformational entropies computed from the distributions of the single strand coordinates. Entropies per coordinate multiplied by the simulation temperature T = 300 K are shown. Only the lesion (or the two adenines facing it) and the two adjacent nucleobase are considered.

These observations are further confirmed by standard deviations of the individual single strand coordinates over a larger portion of the DNA around the lesion (Figures S9-S12). In particular, the undamaged strand facing the solvent at the IN sites is more flexible than the corresponding strand of the control undamaged B-DNA in most of the coordinates (Figure S10). For coordinates related to bending (tilt, roll) and vertical base separation (rise), the high flexibility region extends up to three bases downstream from the damaged site.

The structural details of the N-DNA conformations around the lesion are shown in Figure 5. The conformational states differ in the geometrical arrangement of the 64-PP lesion and, in one case (state 2 of OUT2), the DNA base pairing around the lesion is different from the standard one, with a mispair of the two thymines at the 3’ side of the lesion and involving one of the adenines facing the damage, leaving another adenine (A7) unpaired.

**Figure 5.**
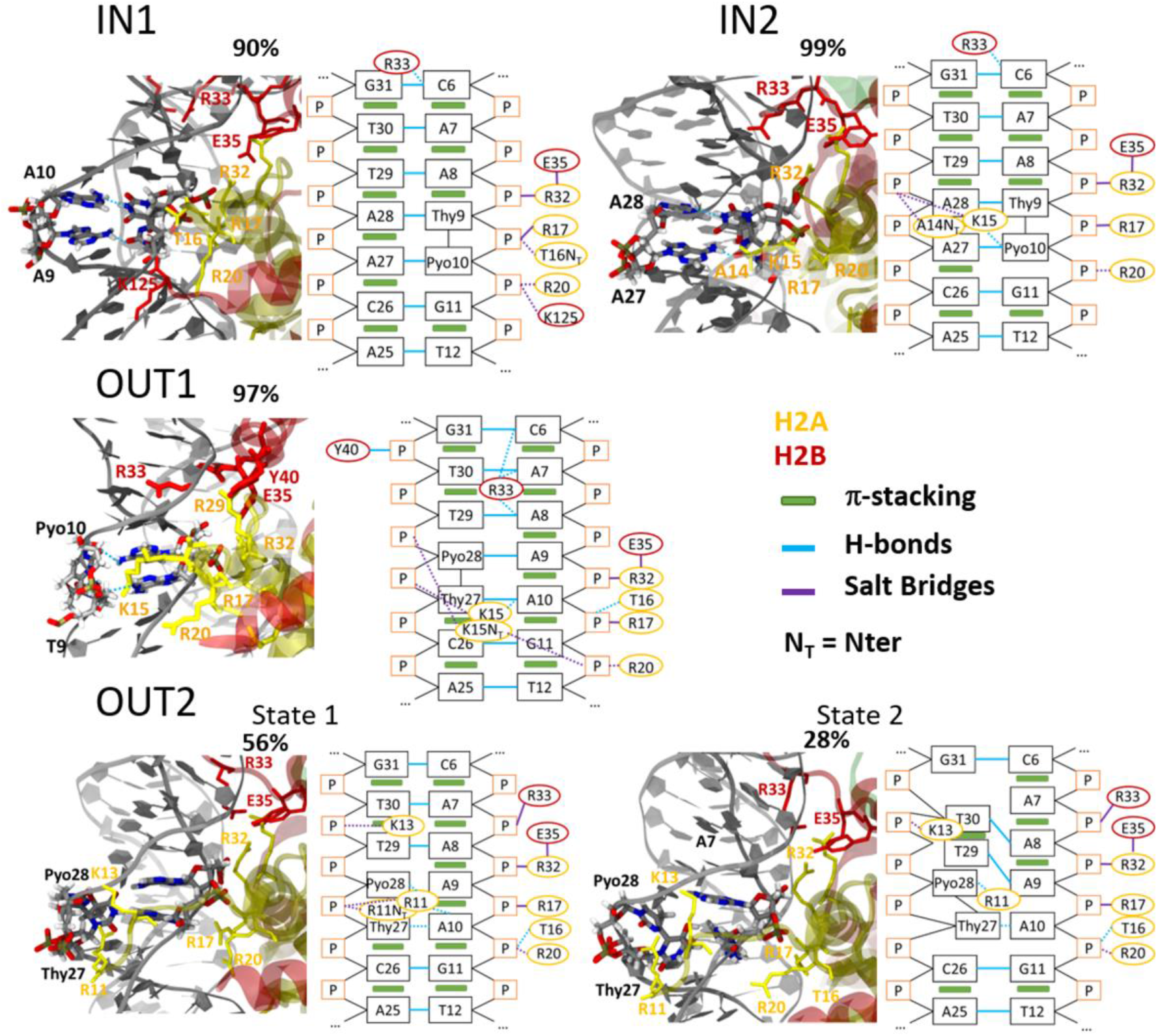
Cartoon representation showing 64-PP at sites 1 and 2 in positions IN and OUT and the associated scheme of the interactions (hydrogen bonds, salt bridges and π-stacking). Solid lines represent stable interactions and dashed lines the transient ones. Residues from the histone are represented in red (H2B) and yellow (H2A). Details of the hydrogen bonding are in Figure S13.

A cooperative network of hydrogen bonds (HB) between the damaged thymines and the complementary adenines is formed. The HB and their occupancies are shown in Figure S13. Both IN sites exhibit an analogous HB network with similar occupancies: two standard Watson-Crick HB between the damaged thymine (Thy) and its complementary adenine (although one with just 50-60 % occupancy), one HB between the Pyo oxygen and its complementary adenine and one HB between this same adenine and Thy. For the more complex and polymorphic OUT sites, the HB network presents interesting differences according to the specific state. In OUT1, the HBs are well defined (Figure S13), but the interaction involves a non-conventional HB between N6 of adenine 28 and O2 of Thy, a situation in sharp contrast with the conservation of standard HB at the IN sites. OUT2 is even more complex, with a conformation impeding the formation of HB between Pyo and the adenine. Only one relatively stable standard HB (82 % occupancy) and multiple unstable ones are observed in state 1 of OUT2 (Figure S13). The latter observation may also be associated with the relatively high flexibility of this specific state (Figure 4). In contrast, in state 2, despite the mispairing, A10 still forms a stable HB with the O5 of the Thy moiety. Moreover, the highly distorted geometry of 64-PP, characterized by two almost perpendicular units, substantially weakens the π-stacking interactions. For the larger part of the trajectories, the π-stacking between Pyo and the adjacent bases is broken while the one between Thy and adenine is preserved. The complementary strand also better conserves its π-stack conformation. However, in state 2 of OUT2, the rotation of one adenine (A10) disrupts the stacking around this base. The consequent loss of HB and π-stacking interactions is compensated by the formation of a pairing between adenine A9 and thymine T29, which justifies the observed low conformational flexibility (Figure 4).

Finally, the detailed analysis of the interactions of the damaged sites with specific residues of the histone core and truncated tails is also schematized in Figure 5. Unsurprisingly, positively charged amino acids (arginines and lysines) develop favourable electrostatic interactions with the negatively charged backbone of the minor groove. Three arginines of H2A, namely R17, R32 and, to a lesser extent, R20, interact with the backbone of the inner strand around the damage. Residues from the H2A N-terminal tails can also interact with 64-PP or the complementary strand as depicted in Figure. These interactions depend on the length of the tail and are not persistent through the full simulation. The most peculiar case is the interaction between R11 and OUT2 where the arginine reaches deeply into the minor groove. In state 1, R11 can interact with both Thy and Pyo fragments as well as their backbone and with A10; while only the interactions with Pyo is allowed in state 2. These differential interactions can contribute to the conformational changes observed for the OUT2 site and justify the different entropies of the two states (Figure 5).

## DISCUSSION AND CONCLUSION

The presence of 64-PP lesions in naked B-DNA induces a considerable increase of the duplex flexibility and may lead to conformational polymorphism. Furthermore, a coexistence of intra-and extra-helical positions for the damage has been reported in previous studies. However, the interaction with compaction proteins, and notably histones, reshapes the free energy landscape of the DNA duplex, significantly altering its flexibility and the accessible conformational space. This is crucial for the nucleotide excision repair initiation protein UV-DDB, which is able to recognize, and bind to, the 64-PP photoproduct within the DNA even if the lesion is facing the nucleosome core and is therefore not directly accessible (25). Until recently, the molecular details of the recognition mechanism have remained elusive, partially also because the crystal structures of nucleosomes containing 64-PP facing the histone core (the IN position) or the solvent (OUT position) are disordered, and hence lack a good resolution, precisely at the damaged sites. The cryo-electron microscopy structures of a 64-PP site detected by DDB in the nucleosome obtained recently by Thomä and coworkers (36) have put forward the hypothesis that, to bind to the buried lesion facing the histone core, UV-DDB changes the translational register of the nucleosome and selectively binds the lesion in an accessible position.

Our MD simulations of nucleosomes containing 64-PP lesions directly facing either the histone core or the solvent, together with the corresponding damaged and control undamaged B-DNA oligomers, provide a complementary picture of the induced structural reorganization and of the consequences for the recognition and repair. The naked, damaged DNA duplex adopts structures that are on average nearly straight, in agreement with the FRET data, but whose flexibility and conformational polymorphism critically depend on the sequence context of the lesion. Nevertheless, the accessible conformational ensemble is wide and the barrier between two specific conformations can be overcome in a few microseconds, as observed for the IN sequence with an intermediate highly bent structure.

The inclusion of the nucleosomal environment significantly changes the above picture, preventing any large fluctuation of the global DNA bending. Besides that, the backbone of the inner strand (damaged or not) is fixed to the histone core by at least three salt bridges involving H2A arginines. When facing the solvent, the DNA damaged strand still experiences an important flexibility and polymorphism of at least three conformational states. Such local deformations should be regarded as signals that can be interpreted and recognized by the repair mechanism, adding to the structural and chemical signal of the lesion itself. When 64-PP is facing the NCP core, the complementary undamaged strand facing the solvent is highly deformed and even more flexible than the corresponding strand in naked undamaged B-DNA. This effect can be rationalized by the fact that the restraints imposed by the histone core restrict the DNA conformational freedom necessary to adapt to the structural distortions induced by 64-PP. Hence, the access to the energetically optimal conformation is limited, a situation that will enhance the local distortion and flexibility around the damage and in particular in the complementary strand facing the solvent. Such observation provides a new and intriguing explanation of the recognition of buried 64-PP by UV-DDB. Indeed, the structural deformation and increased flexibility of the complementary strand will constitute a signal for the presence of the lesion that UV-DDB may interpret, despite the fact that the lesion itself is facing the histone core and is therefore not directly exposed.

The participation of proximal positively charged residues from the histone tails can also be observed on the timescale of our simulations. They can be associated to the development of rather persistent states, and most probably to the partial reshaping of the conformational space of the DNA. The role of the histone tails in the recognition of different lesions has been recently highlighted (37–40), and may be particularly intriguing given the inherent flexibility of the tails and the fundamental signalling role they play in post-translational modifications.

Our results thus indicate a strong sequence dependence of the structure and dynamics of naked DNA duplexes containing 64-PP and the important structural modifications induced by the interactions within the NCP. In particular, we provide a rationale for the recognition of buried 64-PP lesions by the repair enzymes, based on a structural deformation and increase in the flexibility of the complementary strand. In the following research, we aim to proceed to a more systematic study of the structural modifications induced as a function of the position of the lesion in the nucleosome core and by its interaction with the disordered histone tails.

The reader is invited to watch the visualized MD trajectories:

IN 1: https://youtu.be/ALnn_EUDlaM
IN 2: https://youtu.be/cbcsgKdfJd4
OUT 1: https://youtu.be/zHnnfymFEPU
OUT 2: https://youtu.be/Y5sKxHHi55o
IN-naked: https://youtu.be/-5r-5Vf_2Ts
OUT-naked: https://youtu.be/dN7Olc81Tag

## SUPPLEMENTARY DATA

Supplementary Data are available at NAR Online.

## ACKNOWLEDGEMENT

E.B. is grateful for a PhD fellowship from the French Minister of Higher Education and Research. Support from the Université de Lorraine and CNRS is gratefully acknowledged by A.M.

