## Supplementary information for "Impact of the nucleosome histone core on the structure and dynamics of DNA containing pyrimidine-pyrimidone (6-4) photoproduct"

### S1. Supplementary methods

The crystal structures of Osakabe et al. were used as starting points for the nucleosome simulations, the naked damaged and control undamaged DNA oligomers were built in the canonical B-DNA conformation using the nab module of Amber. Net-neutralizing K+ ions and TIP3P waters were added to fill the octahedral periodic simulation box with a minimum distance of 10 angstroms from the solute to the box walls. The systems were equilibrated by first minimizing the energy (500 steps of steepest descent and 500 steps of conjugated gradients), restraining the solute atoms to their initial positions by harmonic restraints using a 25 kcal.mol$^{-1}$.Å$^{-2}$ force constant. This was followed by heating the system from the initial 100 K to 300 K in a 100 ps NVT molecular dynamics, using the Berendsen thermostat with the default coupling time of 1 ps and 25 kcal.mol$^{-1}$.Å$^{-2}$ restraints on the solute. Further, six rounds of energy minimization, each followed by a 50 ps NVT dynamics at 300 K, were performed, gradually releasing the DNA restrains from 5 kcal.mol$^{-1}$.Å$^{-2}$ through 4, 3, 2, 1 to 0.5 kcal.mol$^{-1}$.Å$^{-2}$. Finally, 50 ps of unrestrained NpT dynamics at 300 K and the pressure of 1 atm, using Berendsen thermostat and barostat with coupling times of 5 ps, was performed. The production runs were performed at temperature 300 K and pressure 1 atm using a 2 fs time step and the SHAKE constraints on hydrogen atoms. Snapshots were recorded every 10 ps.

**S2. Supplemetary table**

**Table S1.** Ensemble averages of interhelical (IH) coordinates. Errors are computed as detailed under Methods.

| **System** | **IH shift (°)** | **IH slide (°)** | **IH rise (°)** | **IH tilt (Å)** | **IH roll (Å)** | **IH twist (Å)** |
|---|---|---|---|---|---|---|
| Crystal | 0.36 | -2.4 | 23.34 | 12.74 | 31.72 | 219.95 |
| IN-naked | -2.10 ± 0.14 | 0.16 ± 1.92 | 19.34 ± 1.19 | -6.52 ± 0.85 | 14.30 ± 10.27 | 190.79 ± 11.63 |
| IN-naked last 6 μs | -0.55 ± 0.04 | -0.89 ± 0.20 | 19.65 ± 0.21 | -12.20 ± 0.04 | -16.04 ± 2.49 | 206.01 ± 1.48 |
| OUT-naked | -0.44 ± 0.01 | 3.17 ± 0.01 | 19.80 ± 0.10 | 4.50 ± 0.24 | 16.34 ± 0.58 | 205.57 ± 0.56 |
| IN1 | 0.04 ± 0.06 | 0.23 ± 0.41 | 19.51 ± 0.18 | 37.93 ± 0.15 | 19.58 ± 2.42 | 212.65 ± 1.59 |
| IN2 | -0.42 ± 0.07 | 1.33 ± 0.67 | 19.69 ± 0.09 | 32.46 ± 1.34 | 12.61 ± 0.59 | 213.97 ± 1.13 |
| OUT1 | -2.94 ± 0.04 | 2.78 ± 0.13 | 20.19 ± 0.09 | 20.42 ± 1.25 | -25.40 ± 1.28 | 217.50 ± 0.48 |
| OUT2 – state 1 | -1.42 ± 0.01 | 0.58 ± 0.43 | 20.41 ± 0.14 | 51.32 ± 0.41 | -3.02 ± 2.46 | 206.11 ± 1.45 |
| OUT2 – state 2 | -3.01 ± 0.08 | 4.58 ± 0.06 | 20.00 ± 0.00 | 56.23 ± 1.12 | -6.42 ± 0.65 | 203.62 ± 0.36 |
| Control IN | 0.10 ± 0.01 | 0.63 ± 0.01 | 19.73 ± 0.00 | -6.27 ± 0.06 | -4.26 ± 0.11 | 205.85 ± 0.09 |
| Control OUT | 0.08 ± 0.01 | 0.44 ± 0.00 | 19.81 ± 0.00 | -5.57 ± 0.09 | -3.99 ± 0.09 | 207.34 ± 0.03 |

**S3. Supplementary figures**

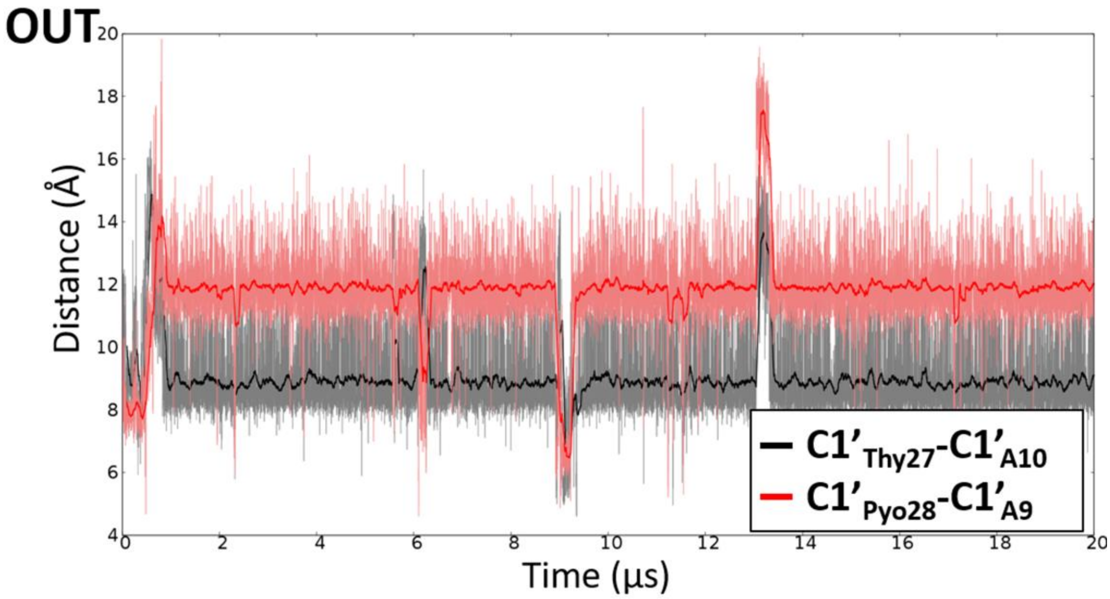


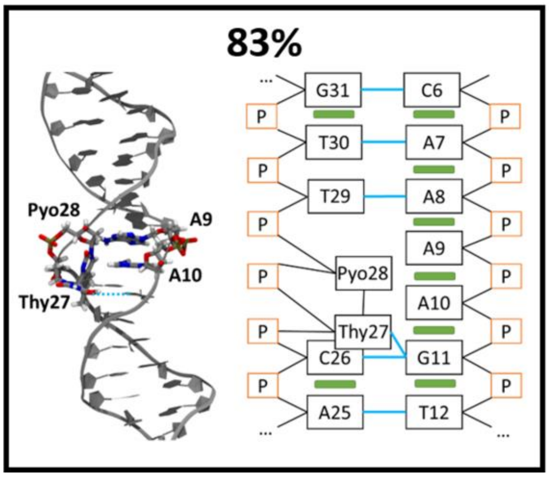


**Figure S1.** Results of the clustering of the MD trajectories for the damaged, naked OUT DNA duplex. The time series of the C1'-C1' distances between the damaged thymines and the opposite adenines defining intra- and extra-helical conformations is also shown. The main cluster is described using a cartoon representation and a scheme for the different interactions (hydrogen bonds in blue solid line and π-stacking indicated by green rectangles).

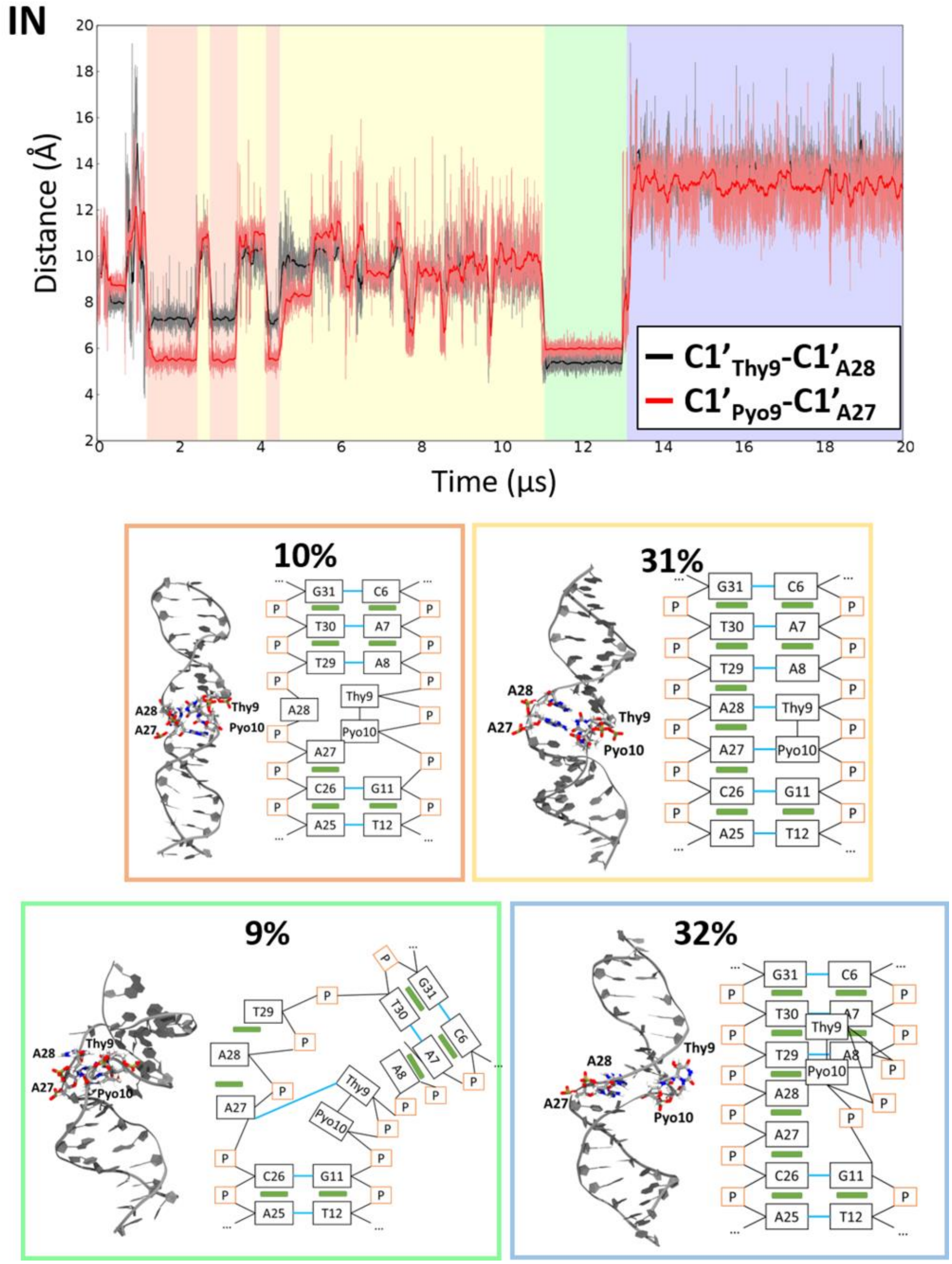


**Figure S2**. Results of the clustering of the MD trajectories for the damaged, naked IN DNA duplex. The time series of the C1'-C1' distances between the damaged thymines and the opposite adenines defining intra- and extra-helical conformations is also plotted. The colours indicate the four different clusters described using a cartoon representation and a scheme for the different interaction (hydrogen bonds in blue solid line and π-stacking in green rectangles).

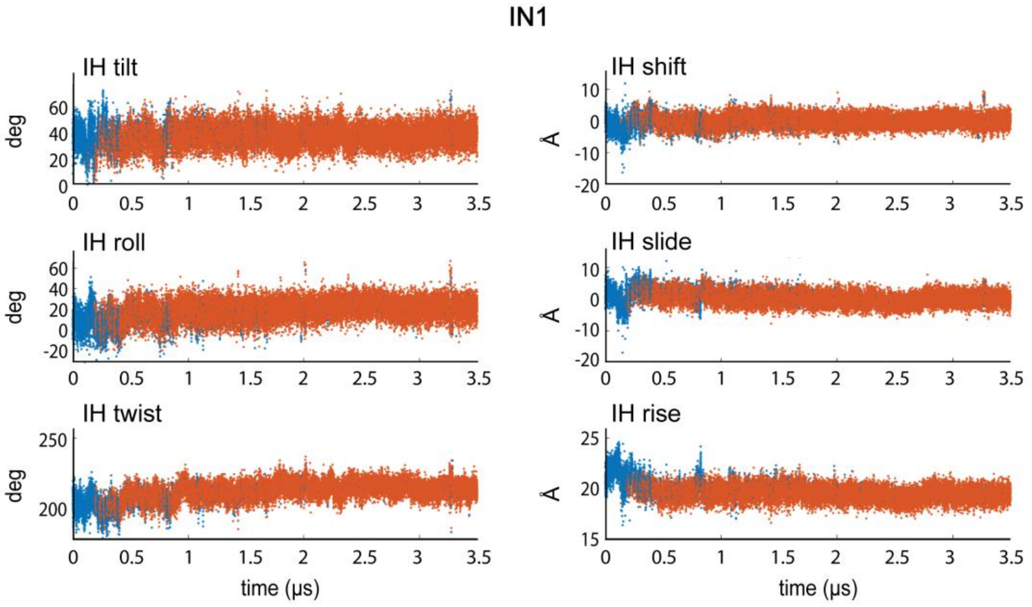


**Figure S3.** Time series of interhelical (IH) coordinates for the IN1 nucleosomal damaged site. The conformational clusters are distinguished by colour. After initial structural rearrangements taking place within the first 0.5 μs (blue), the site adopts one largely dominating conformation (red). The IN2 site behaves very similarly.

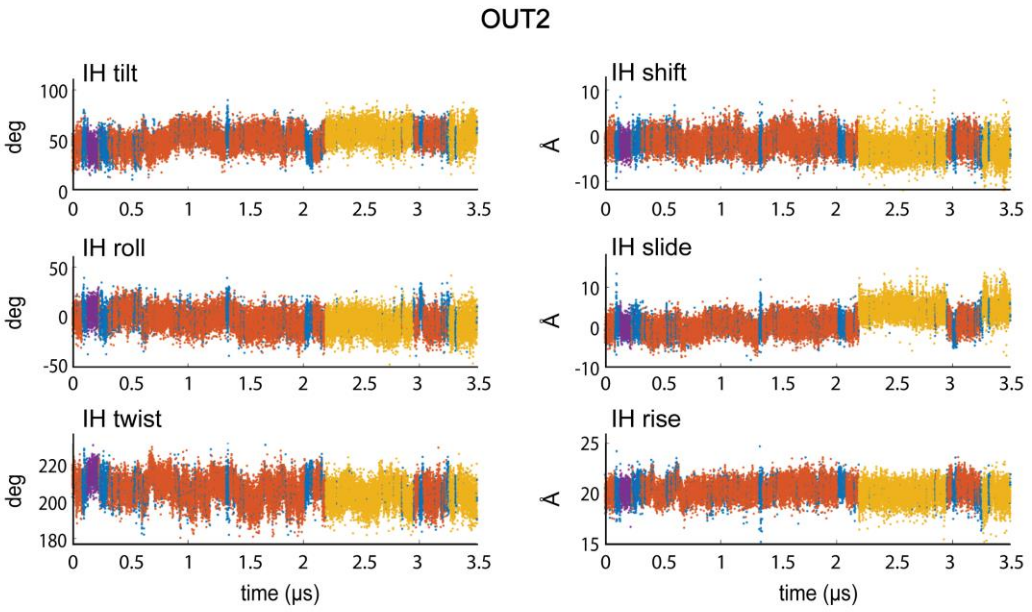


**Figure S4.** Time series of the IH coordinates for the OUT2 nucleosomal damaged site. The conformational clusters are distinguished by colour. Contrary to the IN sites (Figure S3), two major conformational states emerge, denoted as State 1 (red) and State 2 (yellow) in the main text.

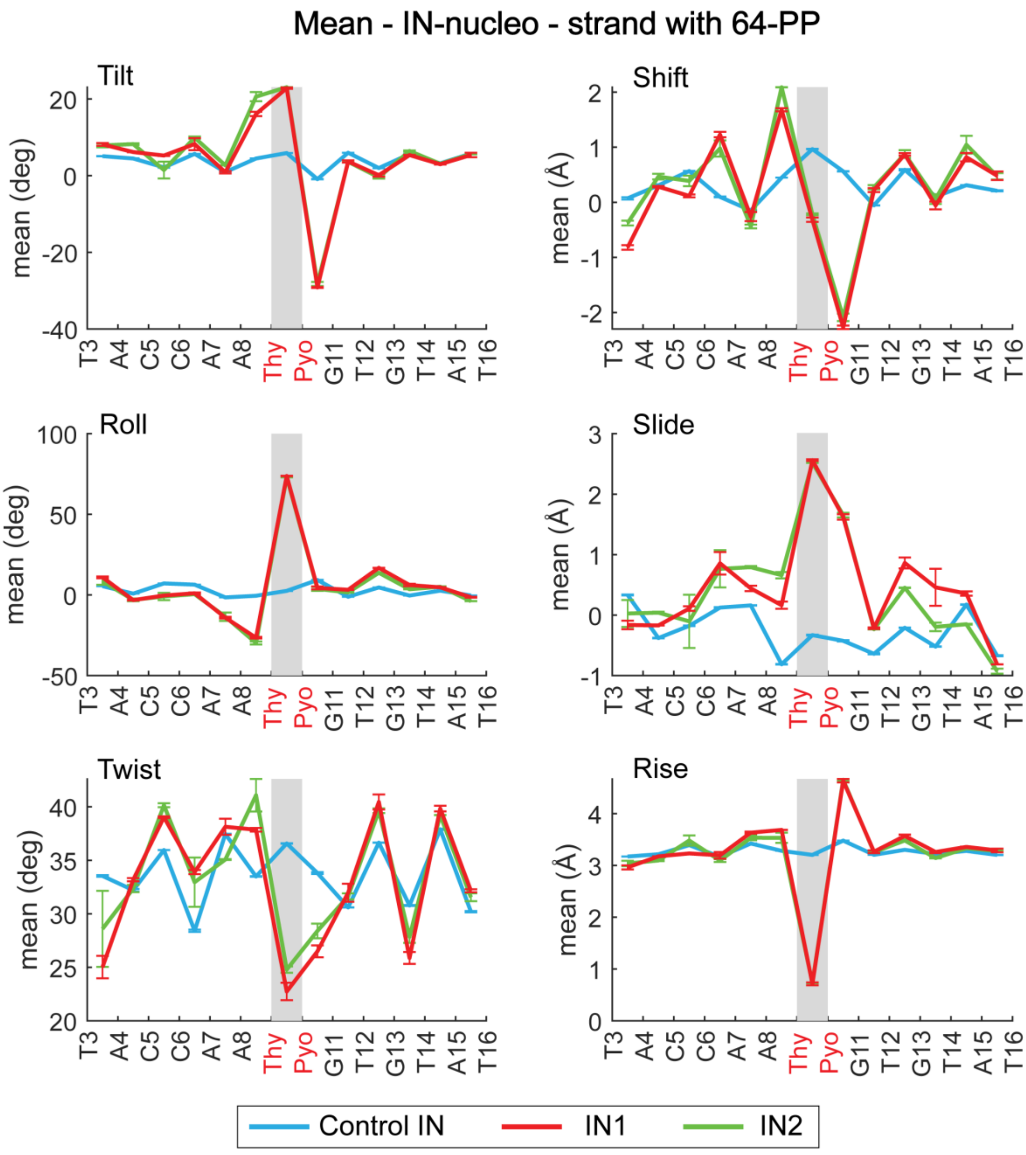


**Figure S5.** Mean values of single strand local coordinates around the IN nucleosomal sites. The strand containing the lesion and facing the nucleosome core is shown. Values of the corresponding strand of the control naked, undamaged DNA is displayed for comparison. The data confirm the near structural equivalence of the two IN sites.

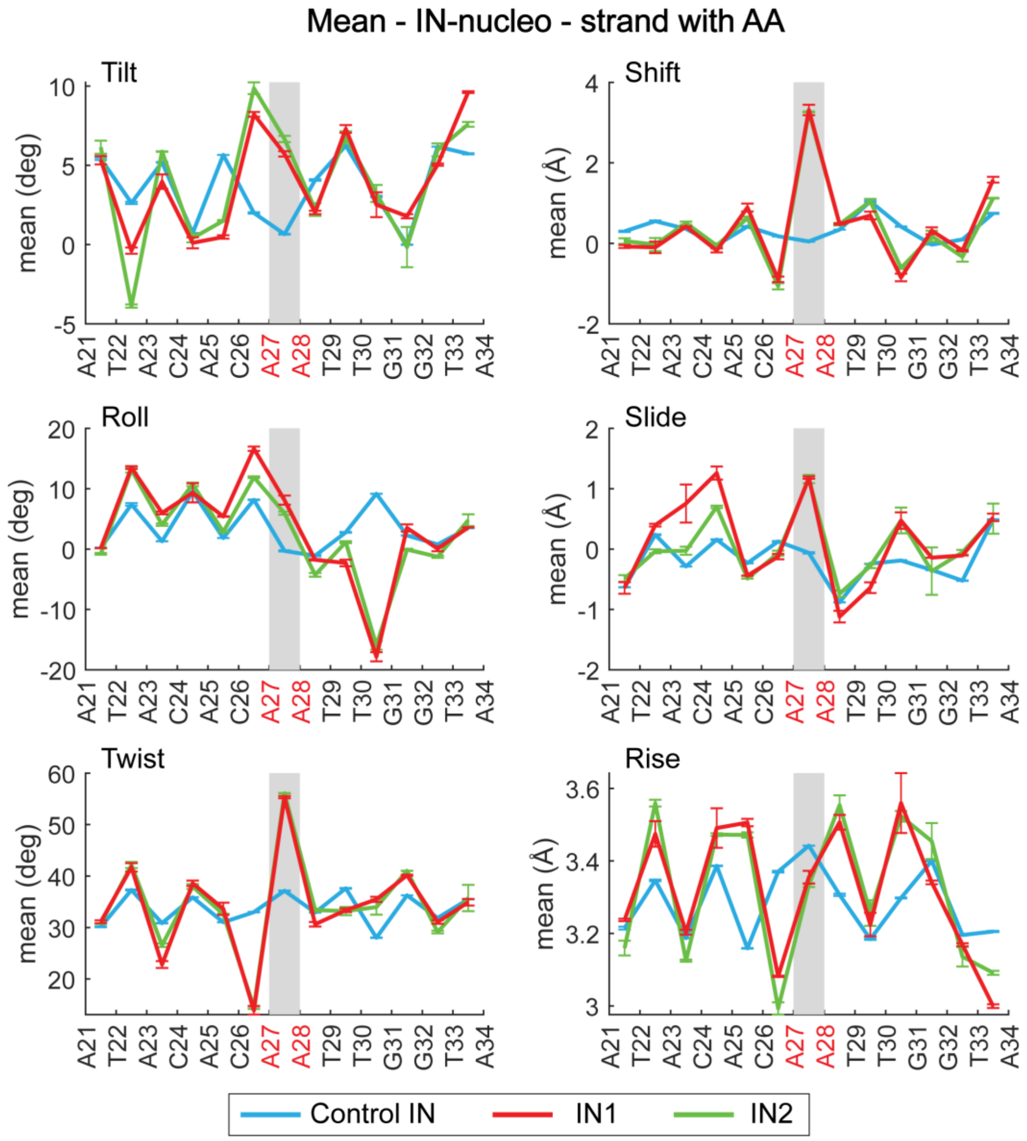


**Figure S6.** Mean values of single strand local coordinates around the IN nucleosomal sites. The strand complementary to the lesion and facing the solvent is shown. Values of the corresponding strand of the control naked, undamaged DNA is displayed for comparison. The data again confirm the near structural equivalence of the two IN sites.

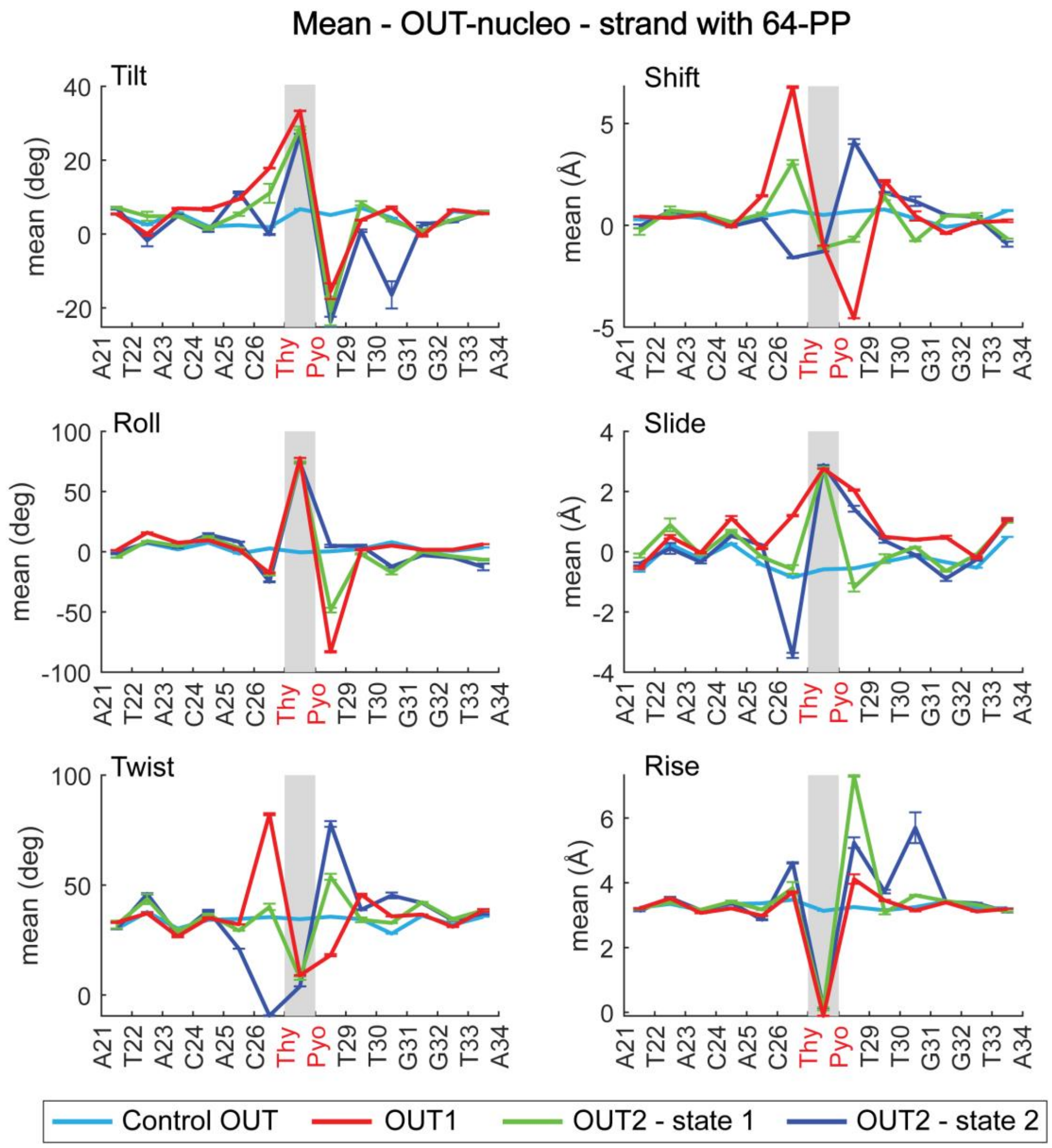


**Figure S7.** Mean values of single strand coordinates for the OUT nucleosomal sites. The strand containing the lesion and facing the solvent is shown. While the OUT1 site adopts just one conformational state, two long-living states are observed at the OUT2 site. Notice the very different geometries of the three states.

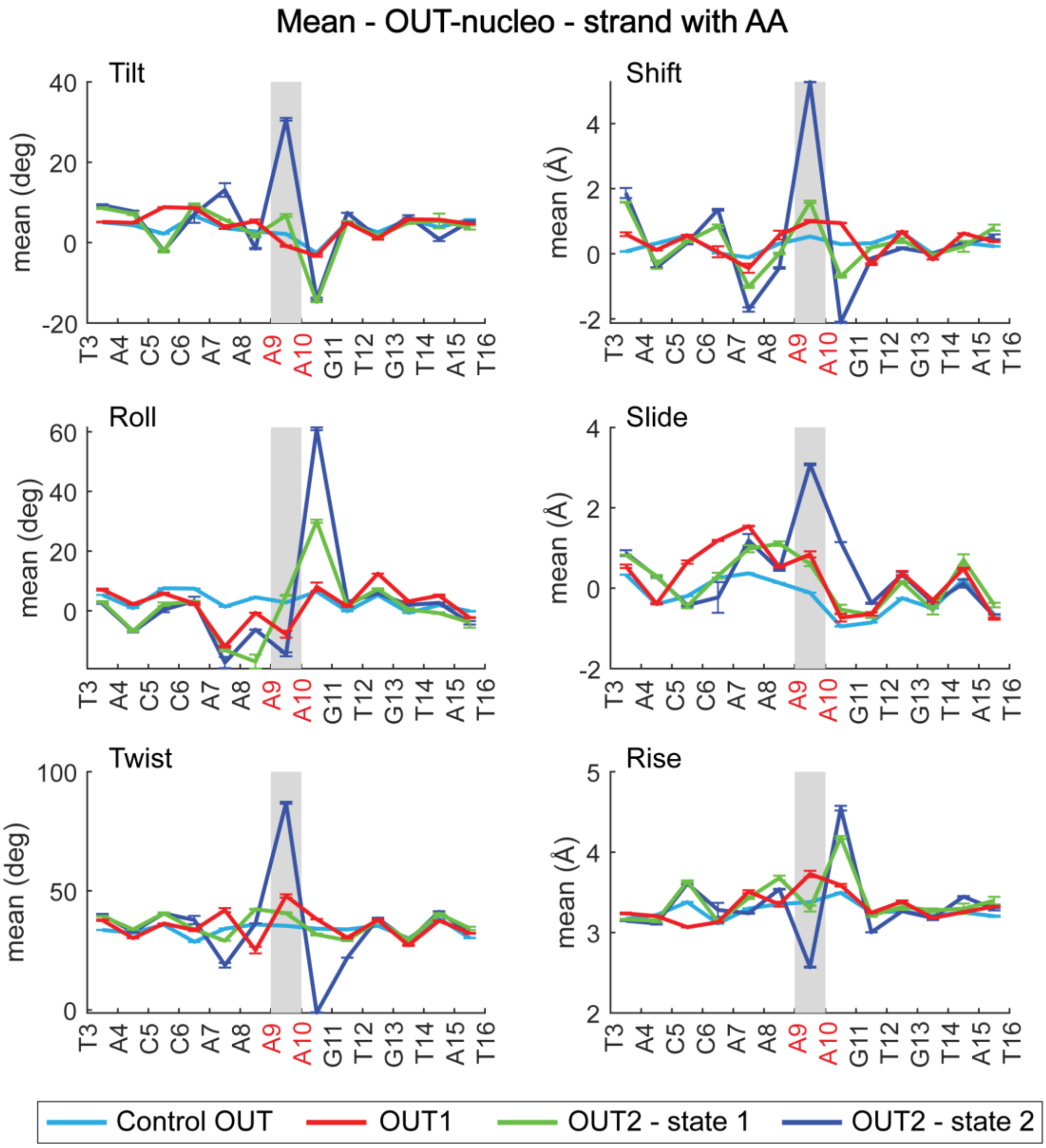


**Figure S8.** Mean values of single strand coordinates for the OUT nucleosomal sites. The strand complementary to the lesion and facing the nucleosome is shown. Once again, the data indicate very different conformations associated with the three OUT states.

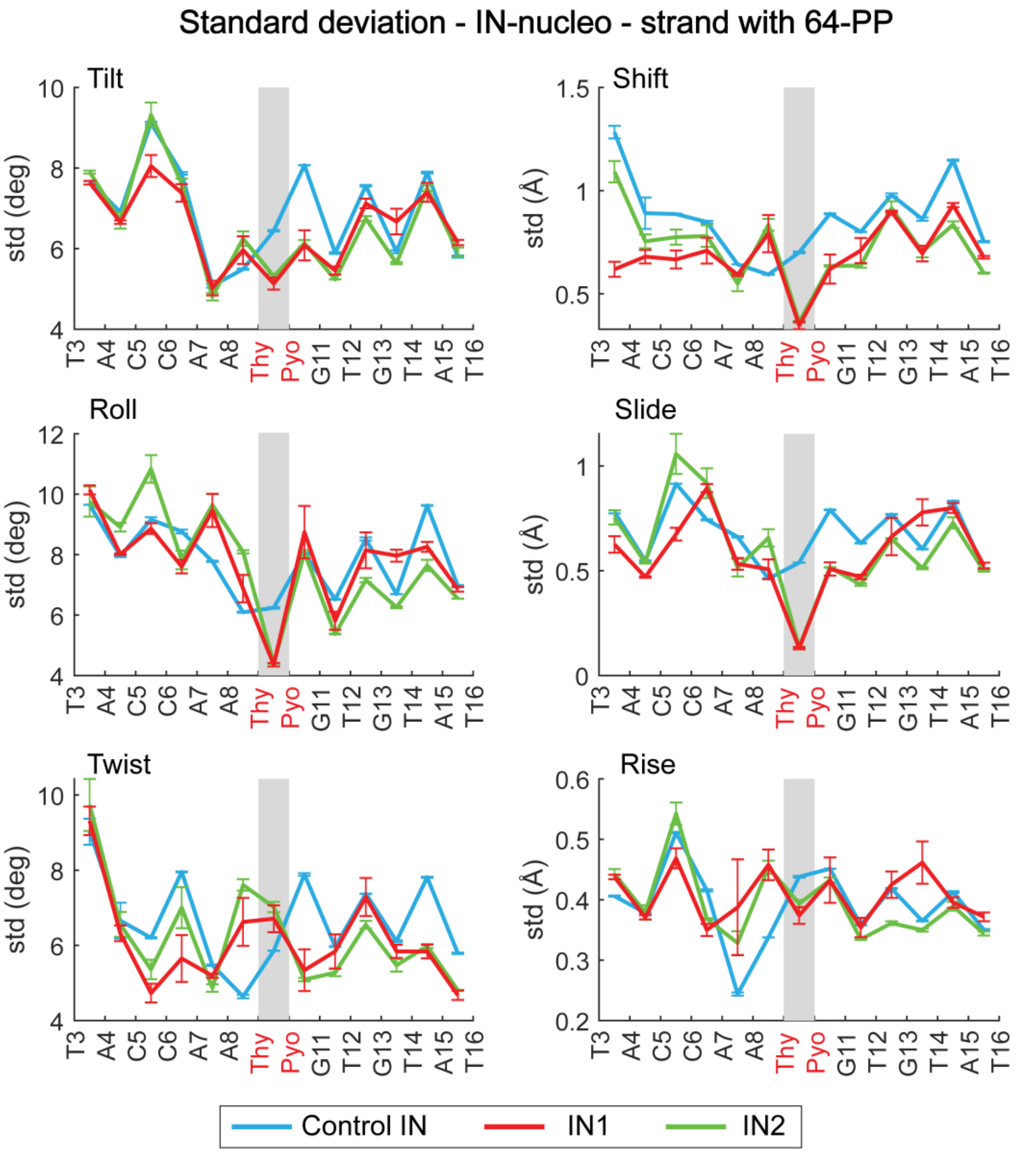


**Figure S9.** Standard deviations of the single-strand local coordinates for the damaged strand at the IN sites. The strand faces the nucleosome core. The corresponding values for the control naked, undamaged B-DNA are also shown.

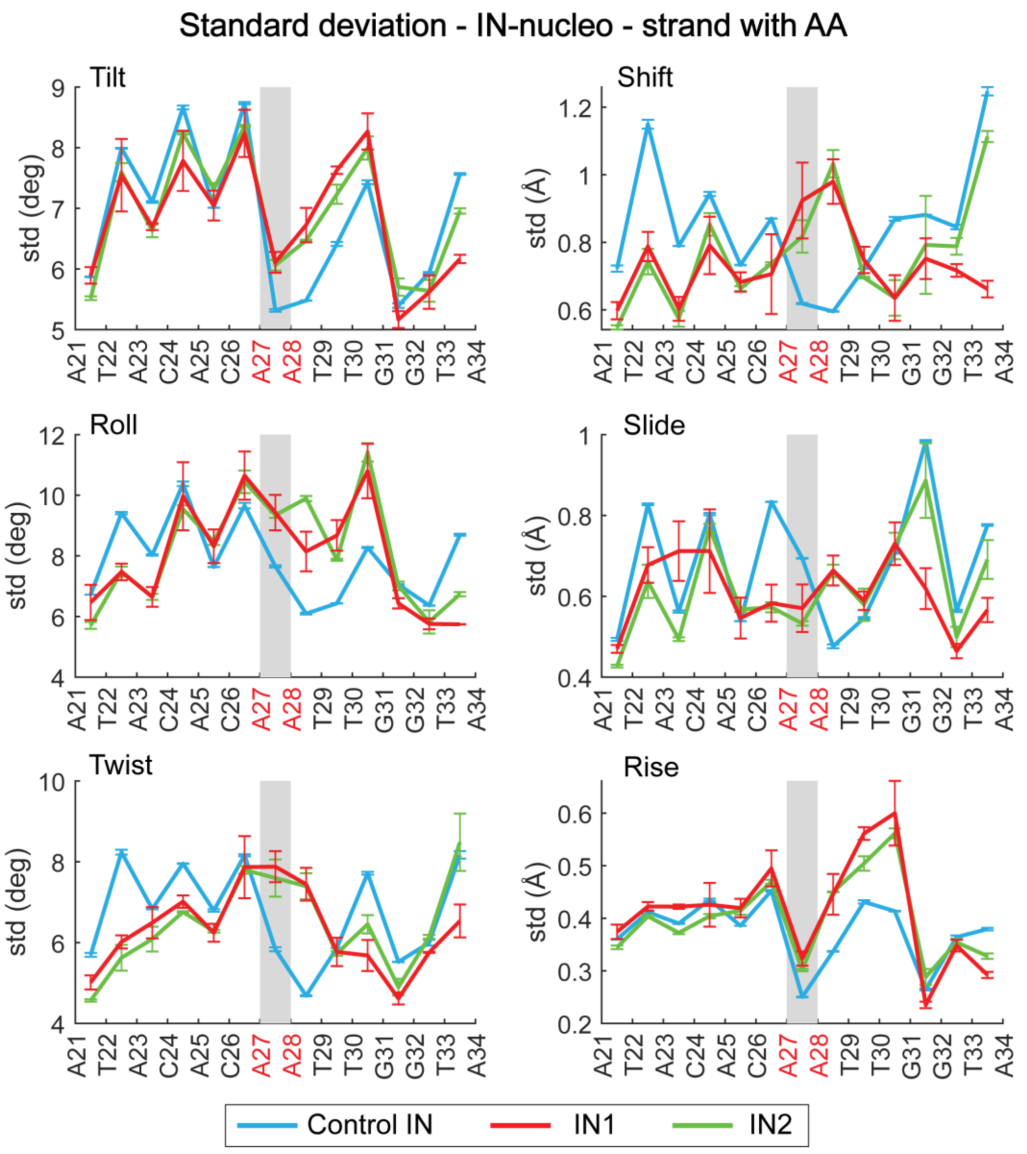


**Figure S10.** Standard deviations of the single-strand local coordinates for the undamaged strand at the IN sites. The strand is exposed to the solvent. The corresponding values for the control naked, undamaged B-DNA are also shown.

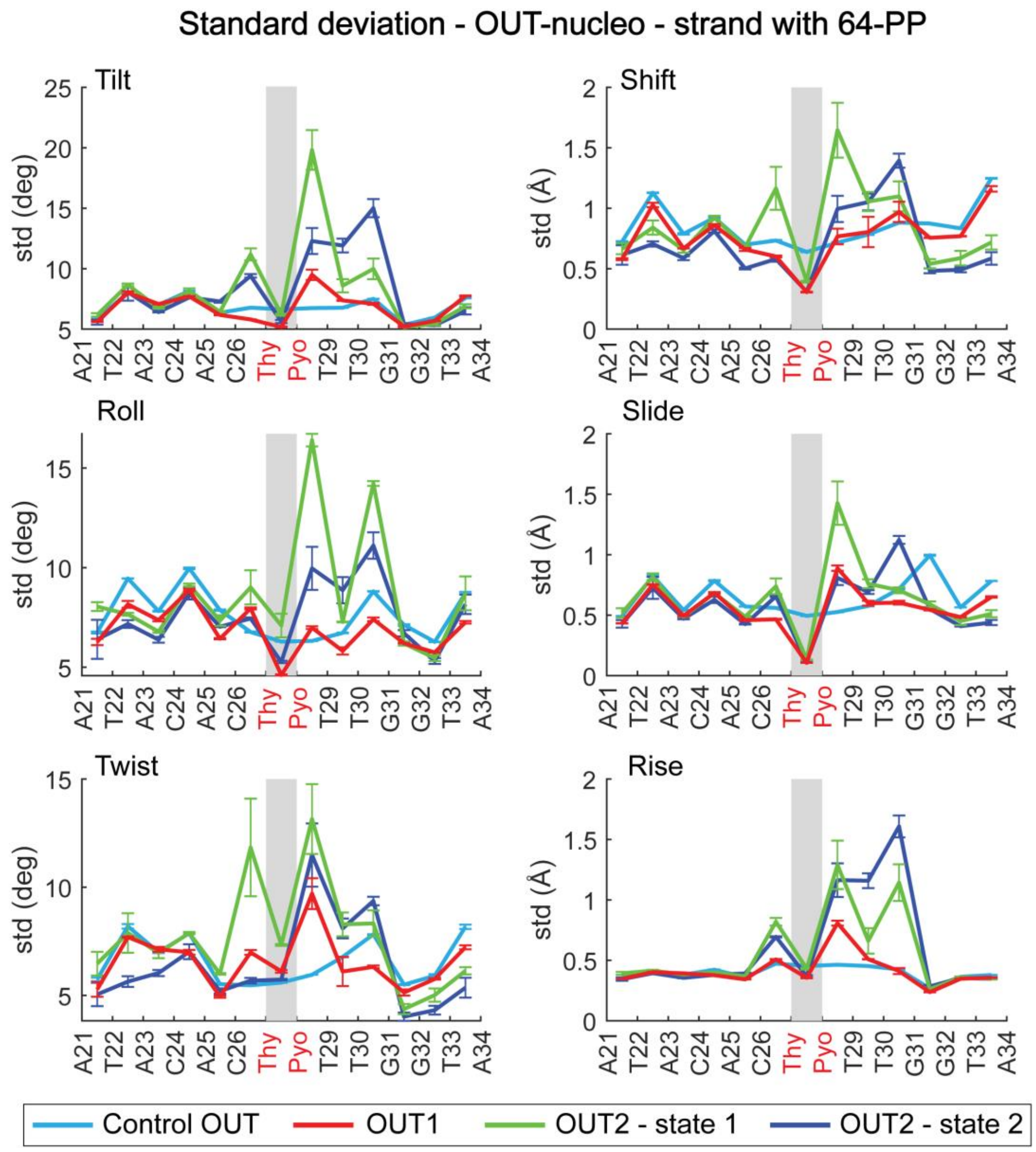


**Figure S11.** Standard deviations of the single-strand local coordinates for the damaged strand at the OUT sites. The strand is exposed to the solvent. The corresponding values for the control naked, undamaged B-DNA are also shown.

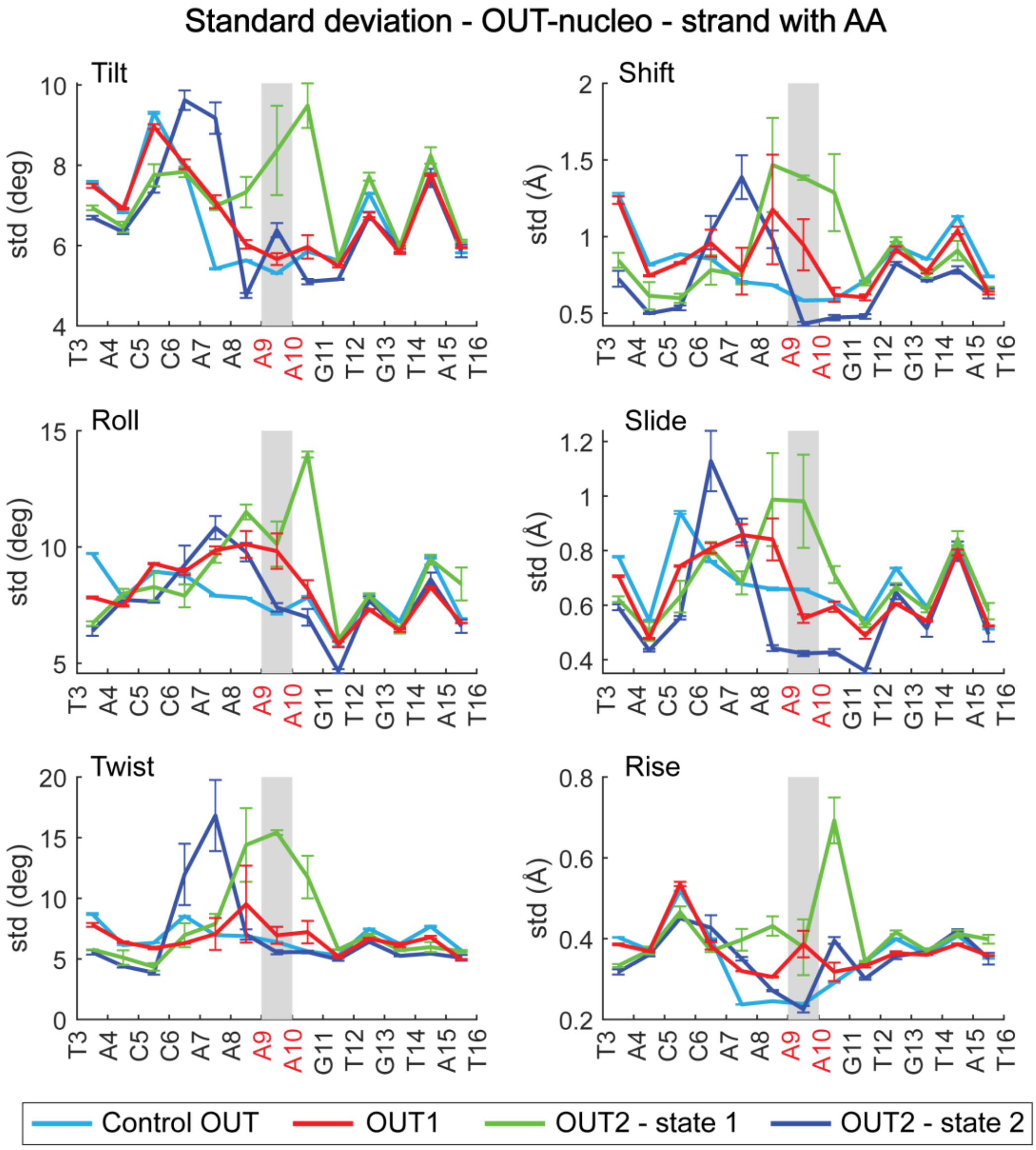


**Figure S12.** Standard deviations of the single-strand local coordinates for the undamaged strand at the OUT sites. The strand faces the histone core. The corresponding values for the control naked, undamaged B-DNA are also shown.

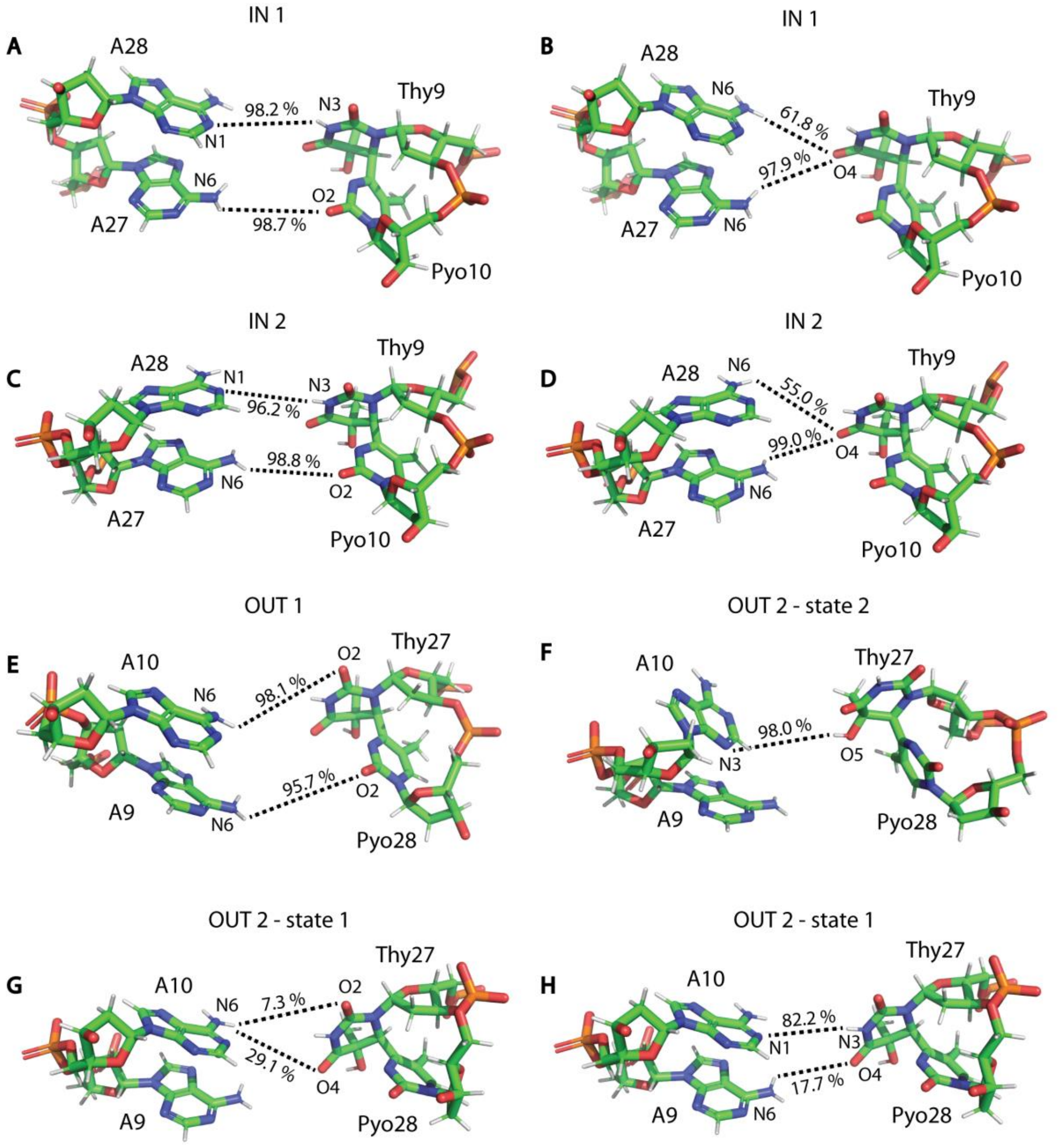


**Figure S13.** Network of hydrogen bonds stabilizing the damaged sites in the nucleosome as inferred from the MD simulations. The bond occupancies are also shown. For clarity, some sites are shown twice with different hydrogen bonds in each case, whereas all these bonds exist simultaneously.
